# Simultaneous estimation of genotype error and uncalled deletion rates in whole genome sequence data

**DOI:** 10.1101/2024.02.06.579174

**Authors:** Nobuaki Masaki, Sharon R. Browning, Brian L. Browning

## Abstract

Genotype data include errors that may influence conclusions reached by downstream statistical analyses. Previous studies have estimated genotype error rates from discrepancies in human pedigree data, such as Mendelian inconsistent genotypes or apparent phase violations. However, uncalled deletions, which generally have not been accounted for in these studies, can lead to biased error rate estimates. In this study, we propose a genotype error model that considers both genotype errors and uncalled deletions when calculating the likelihood of the observed genotypes in parent-offspring trios. Using simulations, we show that when there are uncalled deletions, our model produces genotype error rate estimates that are less biased than estimates from a model that does not account for these deletions. We applied our model to SNVs in 77 sequenced White British parent-offspring trios in the UK Biobank. We use the Akaike information criterion to show that our model fits the data better than a model that does not account for uncalled deletions. We estimate the genotype error rate at SNVs with minor allele frequency > 0.001 in these data to be 3.2 × 10^−4^ (90% CI: [2.8 × 10^−4^, 6.2 × 10^−4^]). We estimate that 77% of the genotype errors at these markers are attributable to uncalled deletions (90% CI: [73%, 88%]).

**Author summary:** A genotype error occurs when the genotype identified through molecular analysis does not match the actual genotype of the individual being analyzed. Because genotype errors can influence downstream statistical results, previous studies have attempted to estimate the rate of genotype errors in a study sample. However, uncalled deletions, which generally have not been accounted for in these studies, can lead to biased error rate estimates. In this study, we formulate a model adjusting for uncalled deletions when estimating genotype error rates. We show that when uncalled deletions are present, this model results in less biased estimates of genotype error rates compared to a model that does not adjust for uncalled deletions. We apply this model to SNVs in 77 sequenced White British parent-offspring trios in the UK Biobank and estimate the genotype error rate and the proportion of genotype errors that are attributable to uncalled deletions at SNVs with minor allele frequency > 0.001.

## Introduction

Genotype errors from high-throughput genotyping technologies can affect the conclusions reached by downstream statistical analysis. For example, genotype errors can reduce the statistical power to detect loci associated with the trait of interest in both pedigree-based linkage studies and case-control genetic association studies [1-3]. Researchers have attempted to mitigate the effects of genotype errors by masking genotypes that are estimated to have a high probability of error prior to statistical analysis [4, 5]. Several methods have also integrated the possibility of genotype errors into linkage analysis to alleviate their effects on the conclusions reached [6, 7].

Given the effects that genotype errors can have on downstream statistical results, researchers may be interested in quantifying the frequency of genotype errors in a particular study. One common approach used to estimate genotype error rates relies on individuals with multiple genotyped samples [8]. However, discordant genotypes will not be observed at a site if these samples all share the same genotype error. Thus, only a lower bound for the genotype error rate can be estimated.

Some genotype errors can be detected by finding Mendelian-inconsistent genotypes in parent-offspring trios. However, only a small percentage of genotype errors at biallelic markers are detectable from Mendelian inconsistencies [9], so probabilistic models have been used in conjunction with the observed number of Mendelian inconsistencies to estimate genotype error rates [10, 11], One limitation of these approaches is that an overall error rate is estimated, rather than error rates that are conditional on the true genotype.

More recent approaches have modeled error rates that depend on both the true genotype and the type of miscall (e.g. the rate at which a major homozygous genotype is miscalled as a heterozygous genotype). Wang et al. use apparent phase violations in three-generation pedigrees to estimate these rates [12], Browning and Browning use a likelihood-based method to estimate the genotype error rates conditional on the true genotype from parent-offspring trio genotypes [13].

No method to our knowledge has adjusted for the effect of uncalled deletions when estimating genotype error rates. An inherited, uncalled deletion creates two genotype errors in a parent-offspring trio. A genotype in a parent with an uncalled deletion will likely be miscalled as homozygous for the non-deleted allele, and the same type of error will be seen in the offspring if the deletion is inherited.

In our study, we model uncalled deletions using an uncalled deletion rate, defined as one half times the probability of an uncalled deletion being present on a randomly chosen genotype in our sample. We develop a model that takes both uncalled deletions and genotype errors into account when calculating the probability of observing a parent-offspring trio genotype, and we use maximum likelihood to simultaneously estimate the uncalled deletion rate and genotype error rates based on observed trio genotypes. Genotype error rates can depend on a marker’s minor allele frequency (MAF), so we count observed trio genotypes within predefined MAF windows and fit a separate model for each MAF window.

We ran two simulation studies to assess model performance. We show that our method results in less biased estimators of genotype error rates if uncalled deletions are present, compared to a model that does not account for uncalled deletions. We also show that our method can accurately estimate the overall genotype error rate and the proportion of genotype errors attributable to uncalled deletions for markers with MAF > 0.001.

Finally, we fit our model to SNV variants in UK Biobank whole genome sequence data for 77 White British trios to obtain estimates and bootstrap confidence intervals for the uncalled deletion and genotype error rates in this dataset. The overall genotype error rate and uncalled deletion rate for SNVs with MAF > 0.001 were estimated to be 3.2 × 10^−4^ (90% CI: [2.8 × 10^−4^, 6.2 × 10^−4^]) and 1.2 × 10^−4^ (90% CI: [1.0 × 10^−4^, 2.7 × 10^−4^]) respectively. We further estimate that 77% (90% CI: [73%, 88%]) of genotype errors at these markers are attributable to uncalled deletions. Using the Akaike information criterion (AIC) [14], we show that our model fits this dataset better than a model that does not take uncalled deletions into account.

## Materials and methods

### UK Biobank sequence data

The UK Biobank release of 200,031 sequenced genomes [15] includes 77 parent-offspring trios whose members have been classified as White British by the UK Biobank [16], We restricted the markers to SNVs having “PASS” in the VCF Filter field, less than 5% missing genotypes, and AAScore > 0.95 [15, 17], and we excluded any SNVs overlapping with a called deletion in any of the 200,031 sequenced genomes. After marker filtering, there were 441,608,608 autosomal SNVs.

### Parental and trio genotypes

If a marker has multiple observed alternate alleles, we combine them into a single alternate allele. We label the major and minor alleles as A and B respectively. Because we restrict our analysis to markers that do not overlap a called deletion, an observed allele is never a deletion. Major allele homozygous, heterozygous, and minor allele homozygous genotypes are denoted AA, AB, and BB respectively. In equations, the AA, AB, and BB genotypes are denoted using the minor allele dose (0, 1, and 2 respectively).

At each marker, we define a trio genotype to be the three genotypes of a parent-offspring trio. We consider the genotypes of the parents to be interchangeable, meaning that two trio genotypes will be considered (and labeled) the same if the father and mother’s genotypes are swapped. Parental genotypes are listed in alphabetical order. For example, AA-AB is a parental genotype in which the two parents have the AA and AB genotypes. Trio genotypes are written as a triplet of genotypes: the two parents’ genotypes are listed first in alphabetical order followed by the offspring’s genotype. As an example, AA-AB-AB represents the trio genotype for which the two parents have the AA and AB genotypes, and the offspring the AB genotype. Table 1 shows the observed trio genotype counts from the 77 UK Biobank White British trios (UKB-WB) [16].

**Table 1.** Autosomal trio genotype counts for 77 UK Biobank White British trios (UKB-WB).

| Trio genotype | Integer representation | UKB-WB | Inconsistent |
| --- | --- | --- | --- |
| AA-AA-AA | 0,0,0 | 33,754,589,673 |  |
| AA-AA-AB | 0,0,1 | 37,896 | X |
| AA-AA-BB | 0,0,2 | 195 | X |
| AA-AB-AA | 0,1,0 | 74,717,572 |  |
| AA-AB-AB | 0,1,1 | 74,518,651 |  |
| AA-AB-BB | 0,1,2 | 34,644 | X |
| AA-BB-AA | 0,2,0 | 37,305 | X |
| AA-BB-AB | 0,2,1 | 23,288,271 |  |
| AA-BB-BB | 0,2,2 | 15,943 | X |
| AB-AB-AA | 1,1,0 | 11,597,232 |  |
| AB-AB-AB | 1,1,1 | 23,436,236 |  |
| AB-AB-BB | 1,1,2 | 11,648,545 |  |
| AB-BB-AA | 1,2,0 | 12,778 | X |
| AB-BB-AB | 1,2,1 | 12,085,809 |  |
| AB-BB-BB | 1,2,2 | 12,100,952 |  |
| BB-BB-AA | 2,2,0 | 74 | X |
| BB-BB-AB | 2,2,1 | 1,985 | X |
| BB-BB-BB | 2,2,2 | 3,927,557 |  |
| Missing | NA | 1,785,549 |  |

Trio genotype counts were from 441,608,608 SNV variants that do not overlap a called deletion. Major allele homozygous, heterozygous, and minor allele homozygous genotypes are denoted AA, AB, and BB respectively. In equations, these genotypes are represented using the integers 0, 1, and 2. The last column indicates whether the trio genotype is Mendelian inconsistent. A trio genotype is considered missing if any member of the trio has a missing genotype.

### Minor allele frequency intervals

We fit our model to the observed trio genotypes from markers whose MAFs are within a specified MAF interval, and we fit a separate model for each interval. This allows model parameters (the genotype error and uncalled deletion rates) to depend on the MAF interval. The 101 MAF intervals are (0, 0.001], (0.001, 0.005], (0.005, 0.01], (0.01, 0.015], …, (0.495, 0.5]. The first two intervals have a length of 0.001 and 0.004, and the remaining intervals have a length of 0.005.

### Overview of the method

Our method assumes that the observed trio genotypes within a MAF interval *I* ⊆ [0, 0.5) are generated under a probabilistic model that incorporates uncalled deletions and miscalled genotypes.

In our model, we start with parental genotypes within MAF interval *I* that have no deleted alleles, which we refer to as the pre-deletion parental genotypes. Deletions are added to a random subset of these genotypes, the frequency of which is determined by an uncalled deletion rate. The genotype of the offspring in each trio is then sampled assuming transmission equilibrium, conditional on the parental genotype.

We assume the trio genotypes in our final sample are drawn proportionally to their true frequencies and that genotype errors occur independently on each genotype during genotyping. Each type of genotype error, defined by the true and observed genotypes, occurs at a fixed rate.

Based on the model described above, we calculate the likelihood of seeing our sample of observed trio genotypes for MAF interval *I* as a function of the uncalled deletion rate and genotype error rates. We use maximum likelihood estimation to obtain an estimate of these parameters. In the following sections, we describe each component of our model in detail.

### Estimating pre-deletion parental genotype frequencies

Pre-deletion parental genotypes are parental genotypes within MAF interval *I* that precede the inclusion of uncalled deletions. There are six types of pre-deletion parental genotypes (AA-AA, AA-AB, AA-BB, AB-AB, AB-BB, BB-BB).

We assume that the frequencies of these parental genotypes (e.g., the frequency of the AA-AB parental genotype) are the same for all markers whose MAFs are within *I*. Because the MAF intervals are narrow, this assumption should be reasonably accurate if the markers are in Hardy-Weinberg equilibrium. We denote the pre-deletion parental genotype frequency for parental genotypes *i, j* in MAF interval *I* as 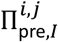 . For example, 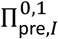 denotes the pre-deletion parental genotype frequency of AA-AB in MAF interval *I*. Because these frequencies are unknown, we estimate 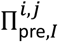 as the observed proportion of parental genotypes *i, j* in MAF interval *I* after excluding Mendelian-inconsistent trios. These estimates are denoted 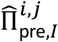.

### Genotypes with uncalled deletions

Deletions are not present in the observed trio genotypes, because we excluded SNVs that overlap with a called deletion. However, we consider the possibility that deleted alleles are present at some markers but are miscalled as the major or minor allele. We label genotypes with a deleted allele as AD or BD, depending on whether the remaining allele is major or minor respectively. We expect deleted alleles to have low frequency at a SNV marker if the called genotypes contain no deleted alleles. Consequently, we do not account for genotypes in which both alleles are deleted because this should rarely occur. In equations, the AD and BD genotypes are denoted using the integers 3 and 4 respectively (recall that the AA, AB, and BB genotypes are denoted using 0, 1, and 2 respectively).

### Estimating true trio genotype frequencies

In our model, the proportion of genotypes with a deleted allele in MAF interval *I* depends on the uncalled deletion rate Γ_*I*_. Each genotype has probability 2Γ_*I*_ of containing a deleted allele, meaning one of the pre-deletion genotype’s alleles (chosen with equal probability) is deleted. This process models the inclusion of uncalled deletions in our sample of trio genotypes.

We estimate the parental genotype frequencies after adjusting for uncalled deletions in each MAF interval, denoted 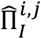, from the previously estimated pre-deletion parental genotype frequencies 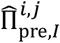 and Γ_*I*_ (see Text S1). We then use the parental genotype frequencies, 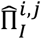, and transmission equilibrium to obtain an estimate of the true trio genotype frequencies 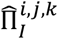, where *k* is the offspring genotype. We will sometimes use the notation 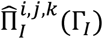 to emphasize the dependency of 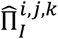 on Γ_*I*_ . Note that the parental genotypes *i, j* in 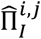 include deleted alleles (i.e., AD and BD). For example, 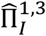 denotes the estimated frequency of the AB-AD parental genotype.

### Modeling genotype errors

In our model, an observed trio genotype is obtained by sampling a trio genotype proportionally to the true trio genotype frequencies 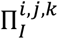, and then introducing genotype errors according to an error model. Let 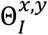 denote the probability that the true genotype *x* ∈ {0,1,2,3,4} is called as the genotype *y* ∈ {0,1,2} at any marker whose MAF is in interval *I*. Genotypes cannot be called as AD or BD because the observed data are SNVs that do not overlap called deletions. We refer to all 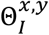 for which *x* ≠ *y* as genotype error rates. The true trio genotype frequencies 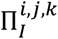 and the genotype error rate parameters 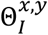 are used to derive the probability of observing a specific trio genotype in MAF interval *I*. We replace each true trio genotype frequency 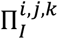 with its estimate 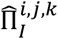, because we do not observe 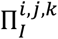 directly.

The rates at which genotypes are called correctly (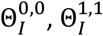 and 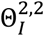), or as homozygous for the non-deleted allele when the true genotype carries a deleted allele (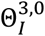 and 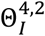), can be expressed in terms of the other ten genotype error rates:

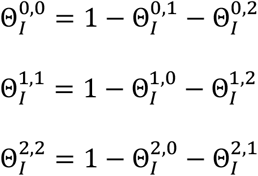

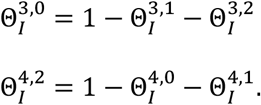

There are 12 error rates in total 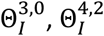, and the ten error rates on the right-hand side of the preceding equations).

In our model, genotype errors occur independently within trio genotypes, meaning that the probability that two genotype errors occur in a single trio genotype is simply the product of the two genotype error rates. Genotype errors also occur independently across different trio genotypes.

To make the optimization problem slightly easier, we assume that 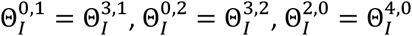, and 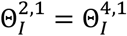 . This means that a genotype with a deletion (AD or BD) will be observed as AA, AB, or BB with the same probabilities as the genotype that is homozygous for the non-deleted allele. This assumption is reasonable because all sequence reads overlapping a marker will carry the same allele when a genotype is homozygous for an allele and when the allele is the non-deleted allele in a genotype carrying a deletion.

So far, we have described the deletion and genotype error processes in our model (see Fig 1 for a graphical summary of these processes). Next, we describe how to calculate the likelihood for observed trio genotypes using this model.

**Fig 1.**
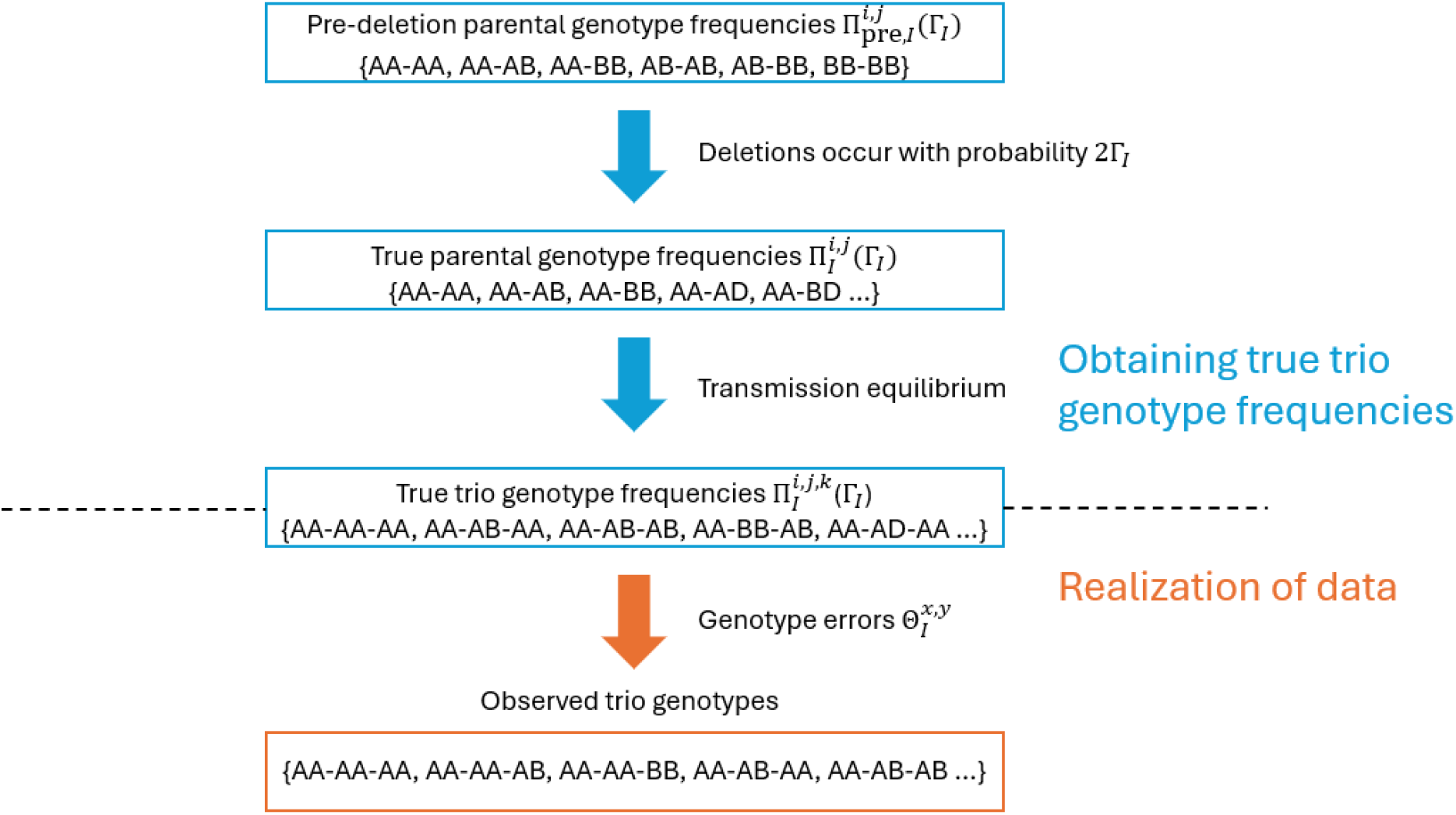
Overview of proposed model. We allow deletions to occur on each pre-deletion genotype with a probability of 2Γ_*I*_. The frequencies of the resulting parental genotypes are referred to as the true parental genotype frequencies. True trio genotype frequencies are calculated assuming transmission equilibrium. The observed trio genotypes are drawn proportionally to these frequencies, and genotype errors are introduced during the sampling process.

### Observed trio genotype probabilities

We can obtain the probability of an observed trio genotype *G*_*m*_ at any marker *m* that falls within MAF interval *I* by summing over all true trio genotypes. For each true trio genotype, we take its frequency as estimated by our model and take the product of this frequency with the genotype error rates corresponding to genotype errors that would lead us to observe *G*_*m*_ (see Equation 1 of Fig 2).

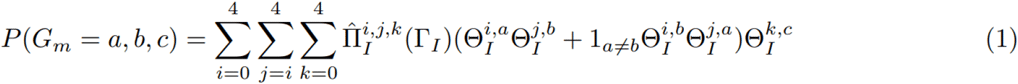

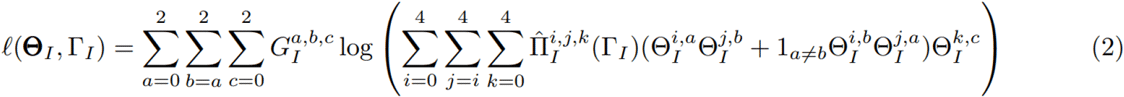

**Fig 2.** Log-likelihood of observed trio genotypes. Equation 1 shows how the probability of a single observed trio genotype is calculated. Equation 2 shows the joint log-likelihood of all the observed trio genotypes in the sample.

Next, we let 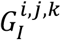 denote the observed count of trio genotype *i, j, k* in *I*. For example, 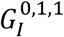 represents the observed count of trio genotype AA-AB-AB in MAF interval *I*. The log-likelihood of observing all trio genotypes within MAF interval *I* is obtained by summing the log-likelihood of observing each trio genotype (see Equation 2 of Fig 2).

### Maximum likelihood estimation

We estimate the genotype error parameters 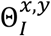 and the uncalled deletion rate Γ_*I*_ by maximizing the log-likelihood *ℓ*(*Θ*_*I*_, Γ_*I*_) (Equation 2 of Fig 2) in the log scale of the parameters using the simulated annealing algorithm (SANN) implemented in the optim() function in R with the default settings [18]. The SANN method produced larger maximized likelihoods than other optimization methods implemented in the optim() function when we fitted our model to the UK Biobank sequence data (see Text S2). Maximum likelihood estimates were exponentiated to their original scales after maximization.

### Estimating the overall genotype error rate and overall uncalled deletion rate

The overall genotype error rate for a set of MAF intervals is the probability that a randomly chosen SNV genotype in the MAF intervals is miscalled. Similarly, the overall uncalled deletion rate for a set of MAF intervals is one half times the probability that a randomly chosen genotype in the MAF intervals contains a deleted allele.

The frequency of genotype *k* ∈ {0,1,2,3,4} in MAF interval *I* can be obtained by summing up the true trio genotype frequencies for which the offspring has genotype *k*. Replacing the true trio genotype frequencies by their estimates, we obtain,

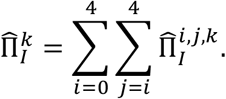

We define the genotype error rate in MAF interval *I*, Δ_*I*_, as the probability that a randomly chosen genotype in MAF interval *I* is miscalled. We estimate Δ_*I*_ using

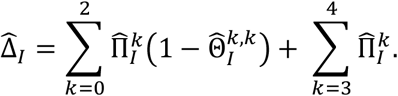

We estimate the overall genotype error rate for a set of MAF intervals as the average of the estimated genotype error rate in each MAF interval, 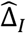, weighted by the number of trio genotypes seen in each MAF interval.

The first MAF interval of (0, 0.001] contains most of the trio genotypes (96.9%) in the UK Biobank sequence data, so including this MAF interval in the weighted average would make the overall genotype error rate for the set of all MAF intervals highly dependent on the genotype error rate in the first MAF interval. Because of this, we report the estimated genotype error rate for (0, 0.001] separately from the estimated overall genotype error rate for the MAF intervals in (0.001, 0.5].

To estimate the overall uncalled deletion rate for a set of MAF intervals, we average the estimated uncalled deletion rates in each MAF interval, 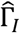, weighting by the number of observed trio genotypes in each MAF interval. We report the estimated overall uncalled deletion rate for the 100 MAF intervals in (0.001, 0.5] and the estimated uncalled deletion rate for (0, 0.001] separately.

Finally, the proportion of genotype errors attributable to deletions for the MAF intervals in (0.001, 0.5] can be estimated by dividing two times the estimated overall uncalled deletion rate for the MAF intervals in (0.001, 0.5] by the estimated overall genotype error rate for the MAF intervals in (0.001, 0.5].

### Confidence intervals

We calculate 95% bootstrap confidence intervals for the uncalled deletion rate and genotype error rates in each MAF interval. To obtain each bootstrap sample, we sample with replacement at the trio level and aggregate the observed trio genotype counts across the sampled trios for each MAF interval. The number of trios in each bootstrap sample is the same as the original number of trios in the dataset. We refit our model on the observed trio genotype counts within each MAF interval in the bootstrap sample and store the resulting estimates of the genotype error rates and uncalled deletion rate.

We repeat this process 100 times to estimate the standard error of each maximum likelihood estimator (corresponding to a genotype error rate or the uncalled deletion rate in the log scale) using the sample standard deviation of the corresponding parameter estimates stored from the 100 iterations. For each parameter, we calculate its 95% bootstrap confidence interval centered around its estimate, assuming a normal distribution for the maximum likelihood estimators in the log scale, and exponentiate the bounds to obtain an interval in the original scale. This prevents the lower bound of our confidence interval from being negative. In the simulation study, this confidence interval had empirical coverage rates closer to 95% compared to a bootstrap confidence interval derived by assuming a normal distribution for the maximum likelihood estimators in the original scale. This assumption may not hold in the original scale because the parameters are rates bounded between 0 and 1.

90% percentile bootstrap confidence intervals for the overall genotype error rate, overall uncalled deletion rate, and proportion of genotype errors attributable to deletions for SNVs with MAF > 0.001 were derived by recalculating these rates for each bootstrap iteration and obtaining the 0.05 and 0.95 quantiles of the resulting bootstrap distributions. We used the percentile method to derive bootstrap confidence intervals for these averaged rates because it was not appropriate to assume that our estimates for these rates followed a normal distribution. The nominal rate of 90% was chosen because of the relatively small number (100) of bootstrap iterations.

### Simulation study 1

We perform a simulation study to calculate the bias of maximum likelihood estimators for the uncalled deletion rate and genotype error rates when observed trio genotypes within a single MAF interval are generated though the deletion and genotype error processes described earlier. We also calculate coverage probabilities of bootstrap confidence intervals for the uncalled deletion rate and genotype error rates. Finally, we compare our estimates of genotype error rates with those obtained from the Browning and Browning method, which does not account for deletions [13].

In the simulation, we generate observed trio genotype counts for four MAF intervals, (0.01, 0.015], (0.05, 0.055], (0.25, 0.255], and (0.49, 0.495], by applying the deletion and genotype error processes with known parameters. These processes are identical to what was described earlier in this section (see Fig 1 for a graphical summary of these processes). To start, we generate pre-deletion genotypes for 100 parental pairs at 3 × 10^8^ markers in each MAF interval, resulting in a total of 3 × 10^10^ pre-deletion parental genotypes per MAF interval.

Minor allele frequencies for the 3 × 10^8^ markers in a MAF interval are drawn from a uniform distribution on the MAF interval. These markers are assumed to be in Hardy-Weinberg equilibrium, and genotypes for parents are drawn based on expected genotype frequencies under random mating.

We use an uncalled deletion rate of 2 × 10^−4^ per allele to add deleted alleles on genotypes. Thus, a deletion occurs on a genotype with a probability of 4 × 10^−4^, in which case a randomly chosen allele is deleted.

We then sample an offspring genotype for each parental pair assuming transmission equilibrium. For example, if a parental genotype is AB-AD, the genotypes AA, AB, AD, and BD occur with equal probability in the offspring at the marker. If by chance, the offspring’s genotype in a true trio genotype is DD, we remove the corresponding trio genotype. This removal maintains the relative frequencies of the true trio genotypes. A DD genotype in the offspring occurs very infrequently because the uncalled deletion rate is low in this simulation study.

We then simulate genotype errors on the true trio genotypes using the genotype error rates in Table 2. These genotype error rates were estimated in an analysis of the UK Biobank data that used a preliminary version of our model. We store the resulting observed trio genotypes.

**Table 2.**
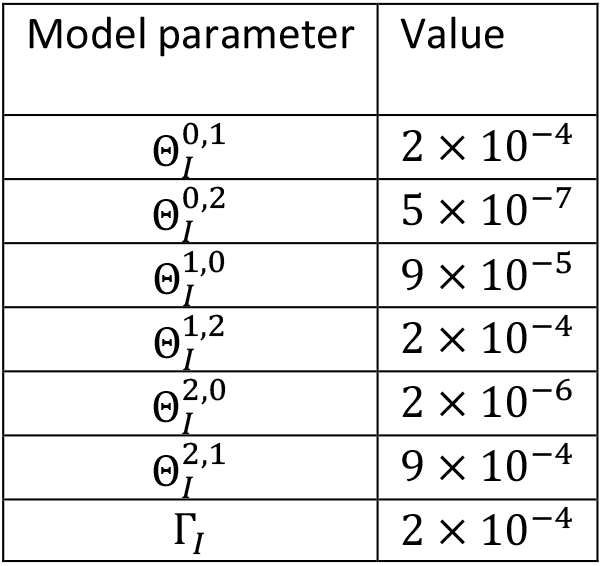
Model parameters used to generate observed trio genotypes in the simulation.

Finally, we fit our likelihood model and the model described in Browning and Browning [13] to estimate genotype error rates (and the uncalled deletion rate when using our model) using the simulated observed trio genotypes (see Maximum likelihood estimation section).

95% bootstrap confidence intervals for genotype error rates (and the uncalled deletion rate when using our model) were generated using the method described in the Confidence intervals section, except we aggregate the observed trio genotype counts across the sampled trios for a single MAF interval, as opposed to for multiple MAF intervals, and refit our model only on this MAF interval. We repeated the above simulation 200 times to calculate the empirical bias of estimators and the coverage probabilities of confidence intervals.

### Simulation study 2

We perform a second simulation to evaluate our model’s estimation of the overall genotype error rate, the overall uncalled deletion rate, and the proportion of genotype errors attributable to uncalled deletions for the MAF intervals in (0.001, 0.5]. We first generate observed trio genotypes for the 100 MAF intervals ((0.001, 0.005], (0.005, 0.01], (0.01, 0.015], …, (0.495, 0.5]).

In each MAF interval, we first generate pre-deletion genotypes for 100 parental pairs. We set the number of simulated markers in each MAF interval so that the observed trio genotype count is similar to that of the corresponding interval in the UK Biobank sequence data. Minor allele frequencies for markers in each MAF interval are drawn from a uniform distribution on the corresponding MAF interval. Genotypes for parents are drawn based on expected genotype frequencies under random mating. Then, we used the same deletion and genotype error processes employed in the previous simulation to generate observed trio genotypes for each of the 100 MAF intervals. The uncalled deletion rate and genotype error rates used are fixed across MAF intervals and are identical to the previous simulation (Table 2).

We obtain the true genotype error rate for each MAF interval, Δ_*I*_, using the true trio genotype counts (before genotype errors are added) and the true genotype error rates. We then average Δ_*I*_ across the 100 MAF intervals, weighting by the trio genotype count in each MAF interval, to calculate the overall genotype error rate for the MAF intervals in (0.001, 0.5].

We obtain the true proportion of genotype errors attributable to deletions for the MAF intervals in (0.001, 0.5] by dividing two times the overall uncalled deletion rate by the overall genotype error rate for the MAF intervals in (0.001, 0.5].

To estimate these quantities, we first fit our likelihood model to the observed trio genotype counts to estimate the uncalled deletion rate and genotype error rates in each MAF interval. Using the process described in “Estimating the overall genotype error rate and overall uncalled deletion rate,” we estimate the overall genotype error rate, the overall uncalled deletion rate, and the proportion of genotype errors attributable to deletions for the MAF intervals in (0.001, 0.5]. We obtain 90% bootstrap confidence intervals for these quantities using the method described in the Confidence intervals section.

Finally, we compare our estimates of the genotype error rate and uncalled deletion rate in each MAF interval, as well as the overall genotype error rate, the overall uncalled deletion rate, and the proportion of genotype errors attributable to deletions for the MAF intervals in (0.001, 0.5] to their true values.

## Results

### Simulation study 1

We show our results from the first simulation study in Table 3. In Figs S1 and S2, we plot estimates for all parameters obtained from the 200 iterations in each of the MAF intervals used to generate the data. The results vary considerably across the different MAF intervals.

**Table 3.**
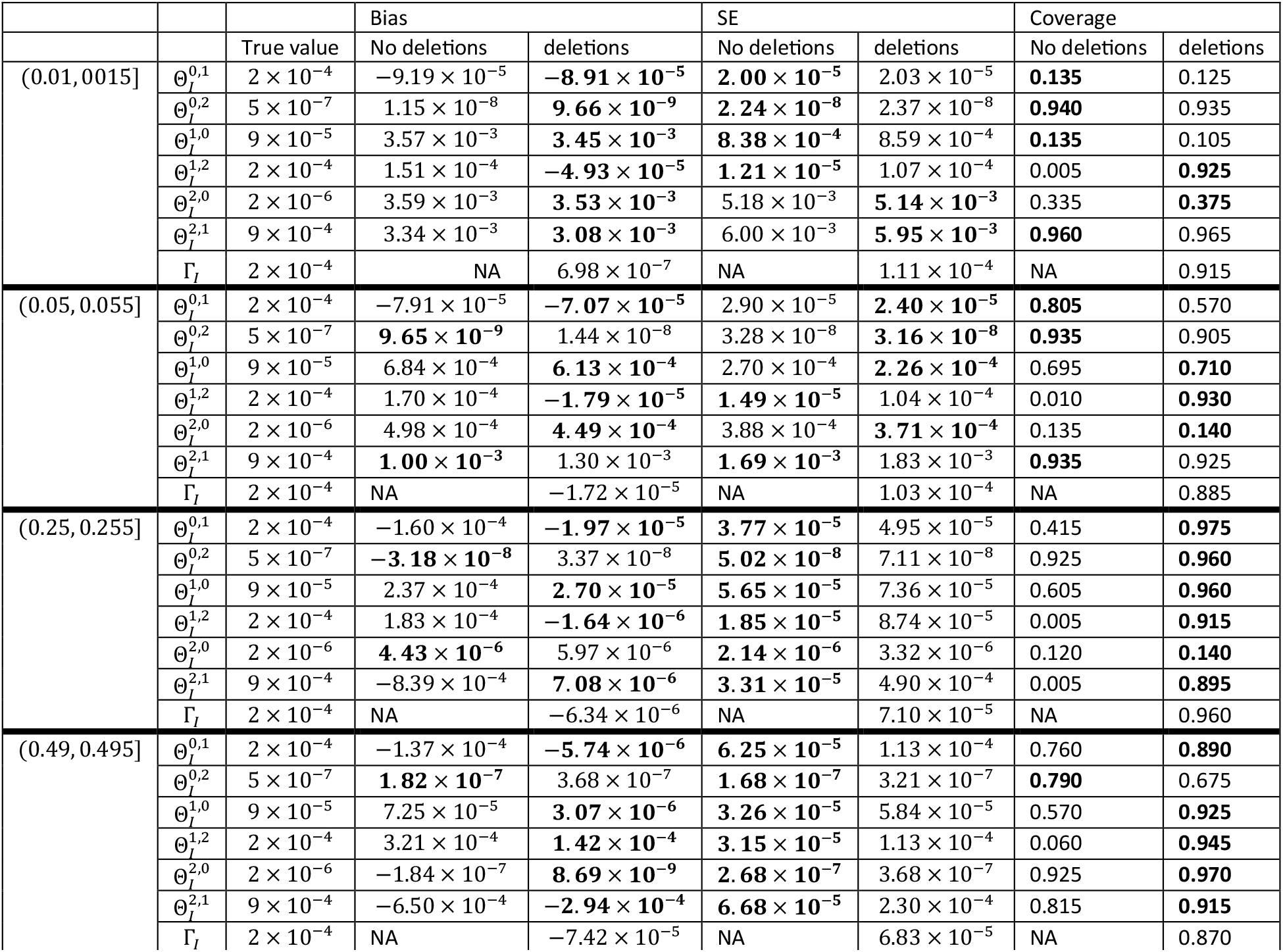
Simulation results.

Genotype error rates were estimated for two models: one with and one without uncalled deletions. The bias of each estimator is calculated by taking the sample mean of each parameter estimate across the 200 simulations and subtracting the true value. The standard error of each estimator is estimated using the sample standard deviation of the parameter estimates across the 200 simulations. The coverage was calculated as the proportion of times the 95% bootstrap confidence intervals captured the true value across the 200 simulations. The bolded values indicate the model that performed better in terms of the corresponding metric (bias, standard error, or coverage).

For the two simulations in which the observed trio genotypes were generated from markers contained in the MAF interval of (0.01, 0.015] or (0.05, 0.055], our model and the model in Browning and Browning [13], which does not account for uncalled deletions, produce similar estimates and bootstrap confidence intervals for all genotype error rates except for 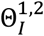 . Our model produces a substantially less biased estimator of 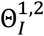. The bootstrap confidence interval generated using our model also captures the true value of 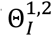at a rate that is much closer to the nominal rate of 0.95. Both models generate accurate estimates for 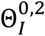, but not for error rates that result in a higher count of observed AA or AB genotypes 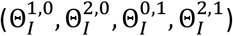. This could be due to an imbalance between genotype counts when these MAF intervals are used (expected BB genotype counts are much smaller than expected AA or AB genotype counts) which make increases in BB genotype counts from genotype errors more apparent than increases in AA or AB genotypes.

For the remaining two simulations in which the observed trio genotypes were generated from markers contained in the MAF interval of (0.25, 0.255] or (0.49, 0.495], our model produces reasonable error rate estimates for the most part, despite some estimators being biased. Overall, our model resulted in estimators for genotype error rates that were less biased but more variable than the Browning and Browning model [13].

Our estimator for the uncalled deletion rate Γ_*I*_ seemed to be effectively unbiased, except when minor allele frequencies for markers were drawn from the highest MAF interval of (0.49, 0.495]. Here, our model tended to underestimate Γ_*I*_.

### Simulation study 2

In Fig 3, we plot the estimated and true genotype error rates for each of the 100 MAF intervals in (0.001, 0.5]. The true overall genotype error rate for the 100 MAF intervals was 6.3 × 10^−4^, while the estimated overall genotype error rate was 6.6 × 10^−4^ (90% CI: [4.2 × 10^−4^, 7.1 × 10^−4^]).

**Fig 3.**
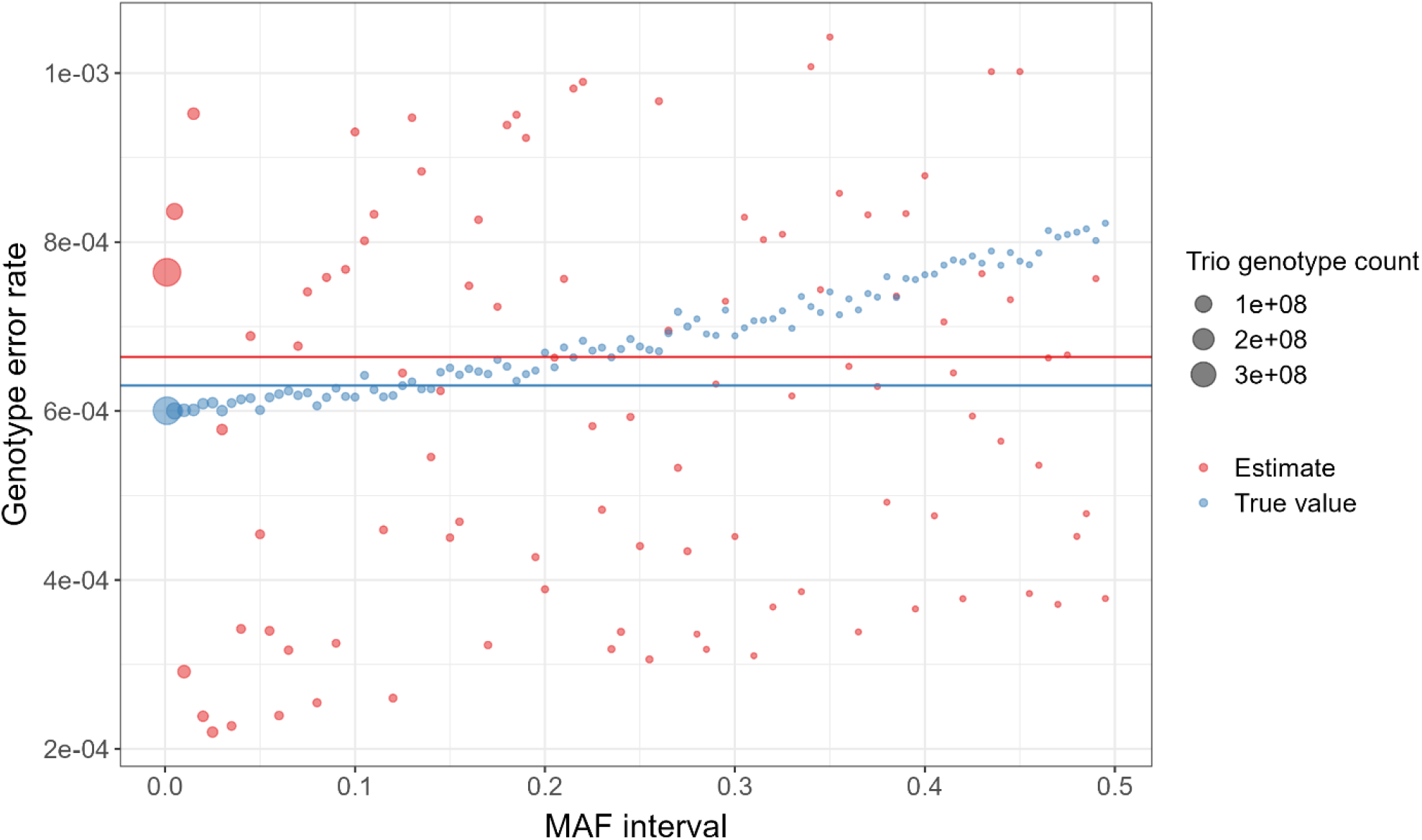
True and estimated genotype error rates for 100 MAF intervals in (0. 001, 0. 5]. The blue and red dots respectively represent the true and estimated genotype error rates for each MAF interval. The size of the dots indicates the number of trio genotypes in the corresponding MAF interval. The blue and red horizontal lines respectively indicate the true and estimated overall genotype error rate for the 100 MAF intervals.

In Fig 4, we plot the estimated uncalled deletion rate for each of the 100 MAF intervals in (0.001, 0.5]. The true overall uncalled deletion rate for the 100 MAF intervals was 2 × 10^−4^, while the estimated overall uncalled deletion rate was 2.2 × 10^−4^ (90% CI: [9.2 × 10^−5^, 2.4 × 10^−4^]) . The true proportion of genotype errors attributable to uncalled deletions was 0.63, while the estimated proportion of genotype errors attributable to uncalled deletions was 0.65 (90% CI: [0.44, 0.68]).

**Fig 4.**
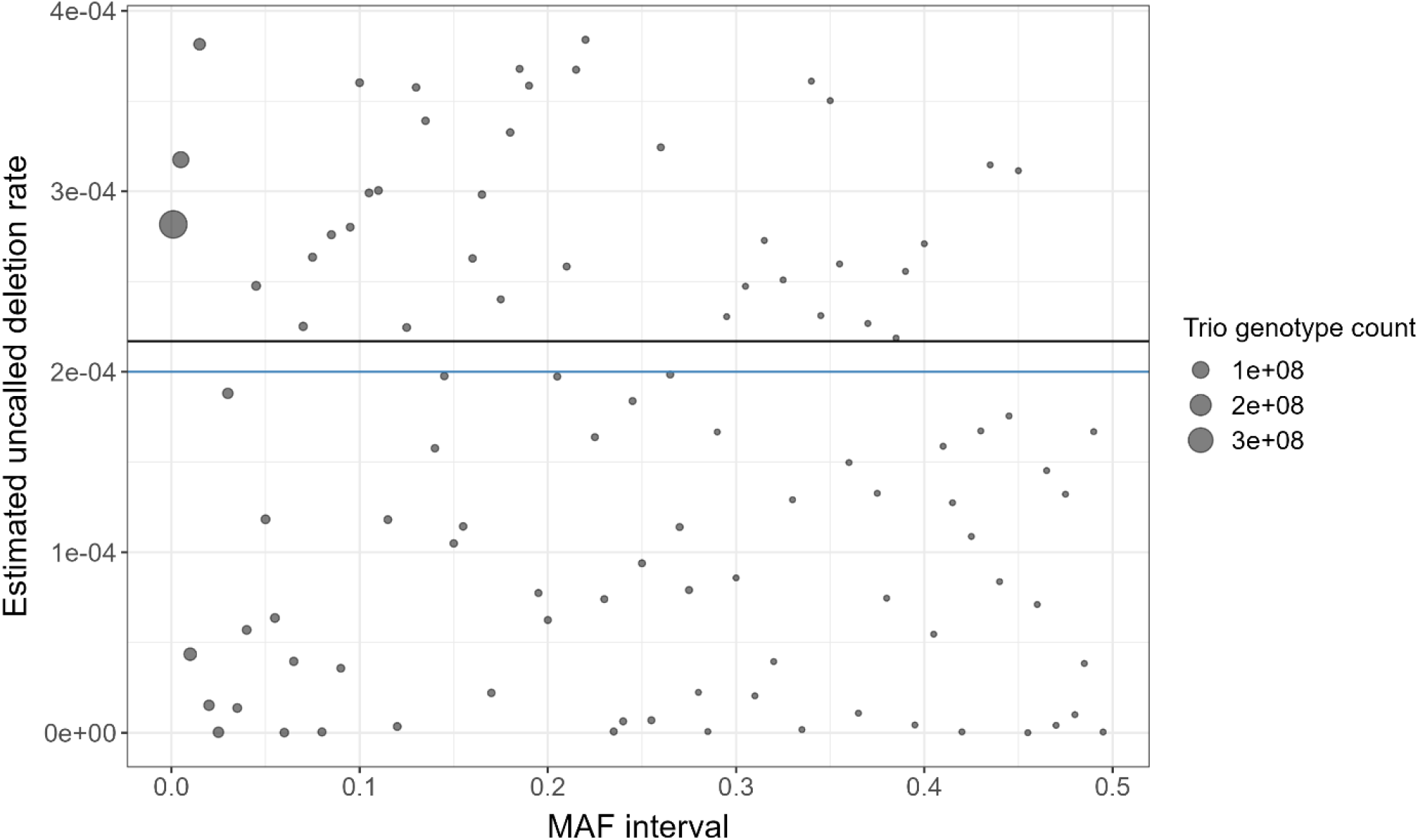
Estimated uncalled deletion rates for 100 MAF intervals in (0. 001, 0. 5]. The dots represent the estimated uncalled deletion rate for each MAF interval. The size of the dots indicates the number of trio genotypes in the corresponding MAF interval. The blue and black horizontal lines respectively indicate the true and estimated overall uncalled deletion rate for the 100 MAF intervals.

### UK Biobank sequence data

Both our model and the Browning and Browning model were fit on each MAF interval in the UK Biobank sequence data. For each model fit, we obtained the parameter estimates, 95% bootstrap confidence interval for each estimated parameter, and AIC of each model [14].

In Fig 5, we plot the estimates and 95% bootstrap confidence intervals from both models. Although bootstrap confidence intervals for parameters calculated using the two models often overlap within the same MAF interval, the estimates between the two models have a noticeably different trend across the lower to higher MAF intervals, except for 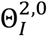 and 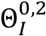 . We see that for the other error rates 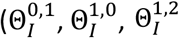 and 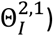, the model with deletions generates higher estimates for error rates that result in a higher count of observed AB genotypes 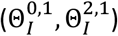 and lower estimates for error rates that result in a higher count of observed AA or BB genotypes 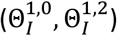.

**Fig 5.**
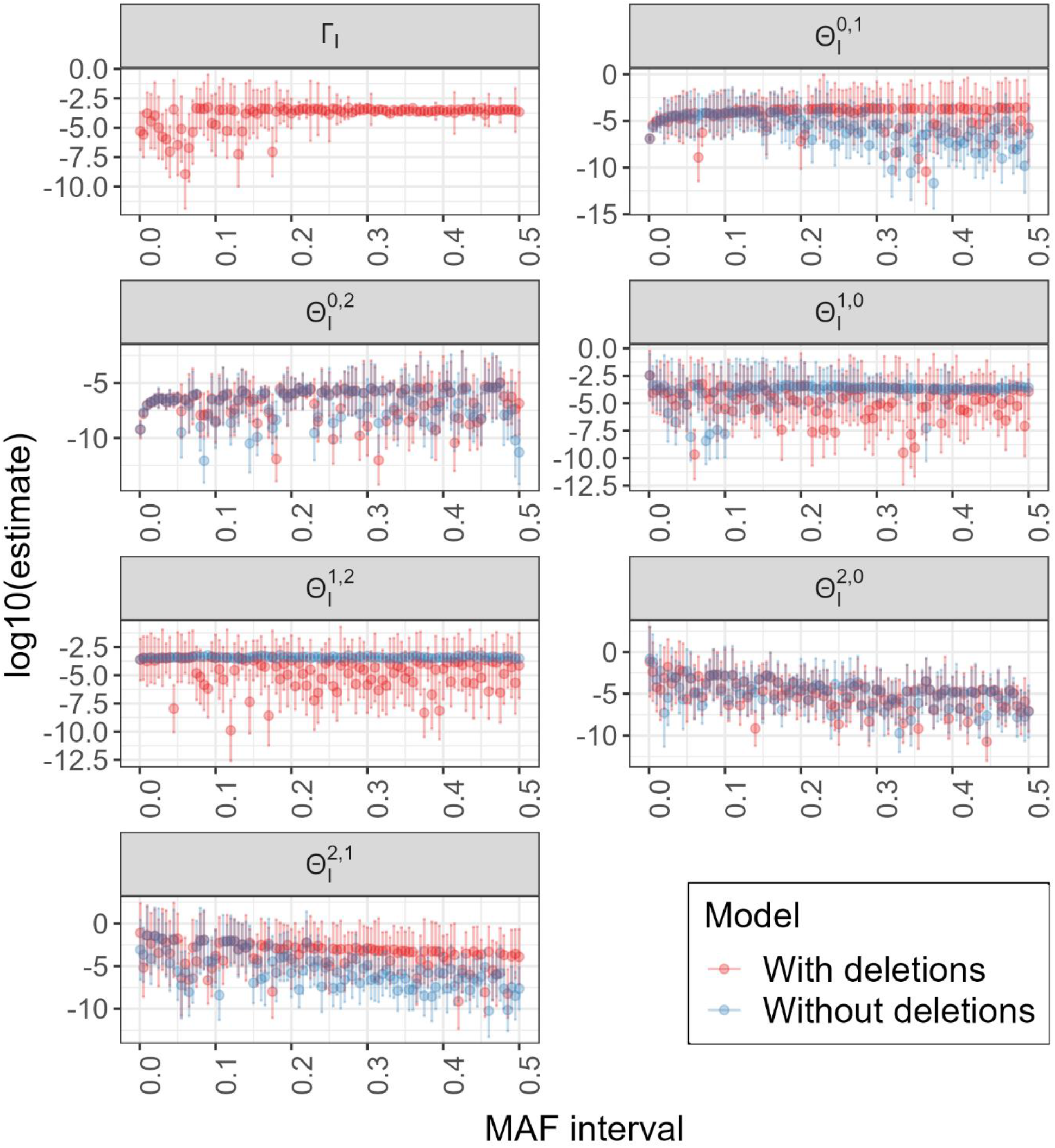
Genotype error rate and uncalled deletion rate estimates and confidence intervals for the UK Biobank sequence data. Genotype error rates were estimated for two models: one with and one without uncalled deletions. Each panel represents a parameter that is being estimated. The x-axis of each panel represents the MAF interval and the y-axis the estimate in the log 10 scale.

Notably, our model estimates an uncalled deletion rate Γ_*I*_ that appears to be largely constant across MAF intervals. This is consistent with our intuition that the rate at which uncalled deletions are present in genotypes should not depend on the MAF of the underlying marker.

We also calculated the AIC of both models fit to each MAF interval. Smaller AIC values indicate better model fit. Compared to the Browning and Browning model [13], our model had a smaller AIC in 81 of the 101 MAF intervals. In Fig 6, we plot absolute differences in AIC between the two models for each MAF interval. We see that AIC slightly prefers the Browning and Browning model at lower MAF intervals.

**Fig 6.**
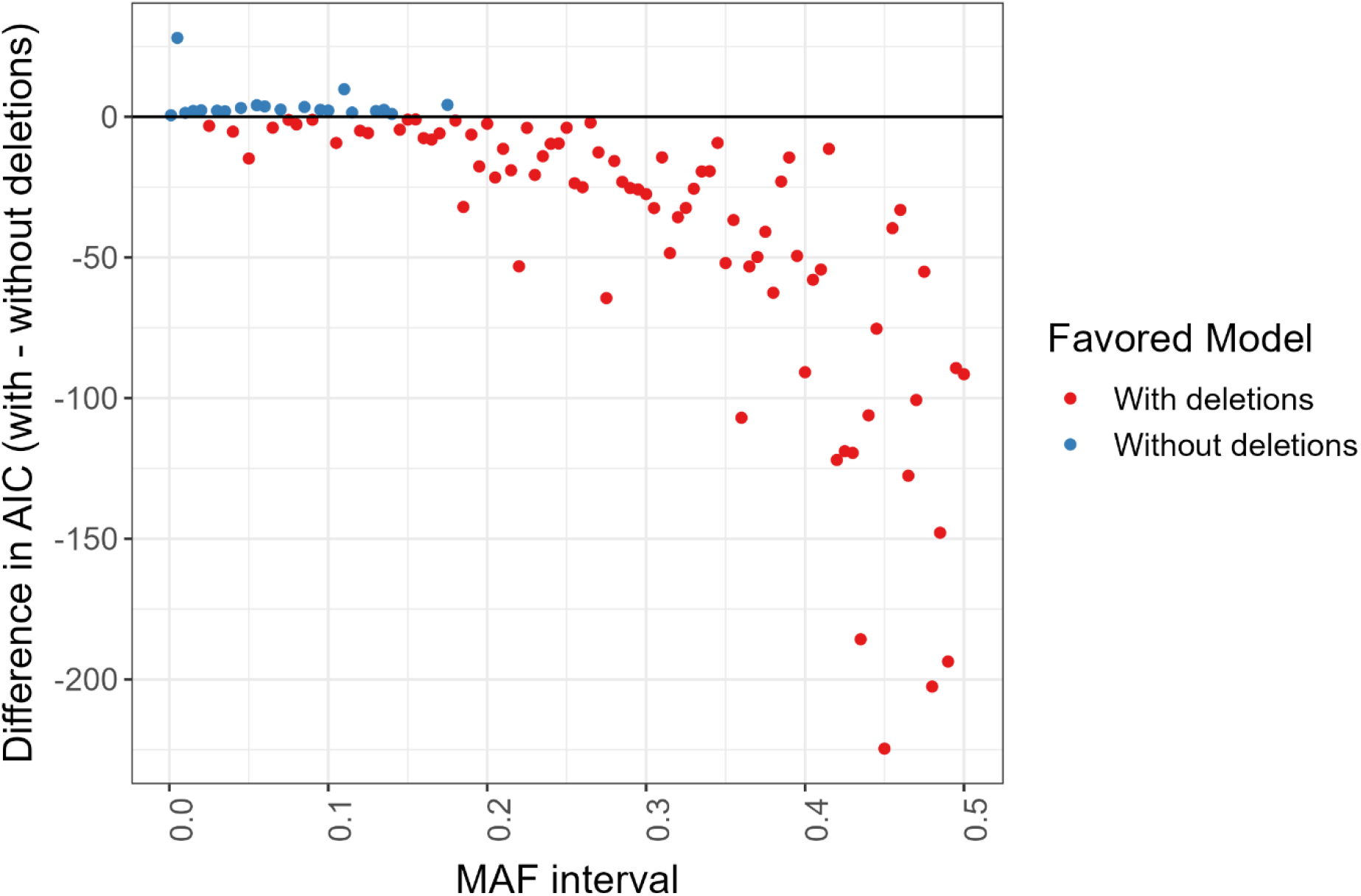
Absolute differences in AIC between the model with and without deletions in the UK Biobank sequence data. The difference in AIC between the models was calculated for each MAF interval in the UK Biobank sequence data. Points below the horizontal line at 0 indicate a preference for the model with deletions, while points above the horizontal line indicate a preference for the model without deletions.

Using our model, the overall genotype error rate and uncalled deletion rate for SNVs with MAF in (0.001, 0.5] were estimated to be 3.2 × 10^−4^ (90% CI: [2.8 × 10^−4^, 6.2 × 10^−4^]) and 1.2 × 10^−4^ (90% CI: [1.0 × 10^−4^, 2.7 × 10^−4^]) respectively. We estimated the proportion of genotype errors attributable to uncalled deletions for SNVs with MAF in (0.001, 0.5] to be 0.77 (90% CI: [0.73, 0.88]). For SNVs with MAF in (0, 0.001], the genotype error and uncalled deletion rate were estimated to be 1.0 × 10^−5^ (90% CI: [1.8 × 10^−6^, 6.8 × 10^−4^]) and 5.1 × 10^−6^ (90% CI: [8.0 × 10^−7^, 3.4 × 10^−4^]) respectively.

## Discussion

Previous studies have attempted to estimate genotype error rates, but no study to our knowledge has controlled for the presence of uncalled deletions in the collected sample, which can lead to biased estimates for these rates. In this study, we present a method for estimating genotype error rates from parent-offspring trio data that allows for the presence of uncalled deleted alleles. Our model allows a proportion of parental genotypes to have uncalled deletions, which can be inherited by the offspring. These uncalled deletions, along with other genotype errors, affect the observed genotypes in our sample. We use a maximum likelihood estimation procedure to estimate the rate of uncalled deletions and genotype errors.

In our simulation studies, our model produced an unbiased estimator for the uncalled deletion rate across a range of marker minor allele frequencies when uncalled deletions are present. Our model also resulted in genotype error rate estimators that were generally less biased than estimators from a model that did not control for uncalled deletions. Finally, our model produced an accurate estimate of the overall genotype error rate, overall uncalled deletion rate, and the proportion of genotype errors attributable to uncalled deletions for a set of MAF intervals.

Using our model, we estimated the overall genotype error rate and uncalled deletion rate at SNVs with MAF in (0.001, 0.5] in the UK Biobank sequence data to be 3.2 × 10^−4^ (90% CI: [2.8 × 10^−4^, 6.2 × 10^−4^]) and 1.2 × 10^−4^ (90% CI: [1.0 × 10^−4^, 2.7 × 10^−4^]) respectively. We further estimated the proportion of genotype errors attributable to uncalled deletions for these SNVs to be 0.77 (90% CI: [0.73, 0.88]). This indicates that uncalled deletions are the primary source of genotype error at SNVs with MAF > 0.001 in these data. Based on the AIC, our model was a better fit to the UK Biobank sequence data than a model that does not account for uncalled deletions.

Our model makes some assumptions that may not be met when fitting the model to real data. First, our model assumes that uncalled deletions occur at a constant rate for every genotype in our sample and that genotyped markers are in Hardy-Weinberg equilibrium. The method could potentially be extended to allow for a more flexible process for introducing uncalled deletions into the sample. Second, we assume that genotypes with deleted alleles (AD or BD) are called as heterozygous (AB), and homozygous for the non-deleted allele at the same rates as genotypes that are homozygous for the non-deleted allele. Relaxing this assumption will increase the number of parameters in the current model, which may require developing a more efficient and robust optimization algorithm.

Additionally, our model is restricted to variants that do not overlap called deletions. Extending the model to incorporate called deletions could improve the estimation of genotype error rates for genotypes with a deleted allele. This would give the added benefit of using genotype data from more markers. However, this would also require us to include the probability of observing a called deletion in the likelihood. Similarly, we could allow for the observation of more than two alleles as an alternative to combining all minor alleles into a composite minor allele.

In this study, we propose a novel approach that uses trio genotype data to simultaneously estimate the rate of uncalled deletions and genotype errors. We show that estimated genotype error rates in UK Biobank sequence data depend on whether uncalled deletions are included in the model, and that the error model that includes uncalled deletions appears to better fit the observed sequence data.

## Acknowledgements

This research has been conducted using the UK Biobank Resource under Application Number 19934. The methodological and analytical work performed in this study was supported by the National Human Genome Research Institute (NHGRI) under award numbers R01 HG008359. The content is solely the responsibility of the authors and does not necessarily represent the official views of the National Institutes of Health or the UK Biobank.

