## Supplemental information for "Simultaneous estimation of genotype error and uncalled deletion rates in whole genome sequence data"

### Text S1: Derivation of $\widehat{\Pi}_I^{i,j}$

Our estimates for the true parental genotype frequencies $\widehat{\Pi}_I^{i,j}$ in MAF interval $I$ are derived using the estimated pre-deletion parental genotype frequencies $\widehat{\Pi}_{\text{pre},I}^{i,j}$ and the uncalled deletion rate $\Gamma_I$. Recall that $\widehat{\Pi}_{\text{pre},I}^{i,j}$ is estimated as the observed proportion of each parental genotype pair in MAF interval $I$ after excluding Mendelian-inconsistent trios.

Note that there are only six possible pre-deletion parental genotype pairs (AA-AA, AA-AB, AA-BB, AB-AB, AB-BB, BB-BB), but there are 15 possible true parental genotype pairs (AA-AA, AA-AB, AA-BB, AA-AD, AA-BD, AB-AB, AB-BB, AB-AD, AB-BD, BB-BB, BB-AD, BB-BD, AD-AD, AD-BD, BD-BD). We estimate $\widehat{\Pi}_I^{i,j}$ by considering the ways in which these 15 genotype pairs can occur through the replacement of one or more alleles with deletions in pre-deletions parental genotype pairs. For example, AA-AD can arise from AA-AA if there is a deletion of any of the four alleles and also from AA-AB if there is a deletion of the B allele. Thus, we have that $\widehat{\Pi}_I^{0,3} = 4\widehat{\Pi}_{\text{pre},I}^{0,0}\Gamma_I(1-2\Gamma_I) + \widehat{\Pi}_{\text{pre},I}^{0,1}\Gamma_I(1-2\Gamma_I)$. We derive all $\widehat{\Pi}_I^{i,j}$ this way:

$$\widehat{\Pi}_I^{0,0} = \widehat{\Pi}_{\text{pre},I}^{0,0}(1-2\Gamma_I)^2$$

$$\widehat{\Pi}_I^{0,1} = \widehat{\Pi}_{\text{pre},I}^{0,1}(1-2\Gamma_I)^2$$

$$\widehat{\Pi}_I^{0,2} = \widehat{\Pi}_{\text{pre},I}^{0,2}(1-2\Gamma_I)^2$$

$$\widehat{\Pi}_I^{0,3} = 4\widehat{\Pi}_{\text{pre},I}^{0,0}\Gamma_I(1-2\Gamma_I) + \widehat{\Pi}_{\text{pre},I}^{0,1}\Gamma_I(1-2\Gamma_I)$$

$$\widehat{\Pi}_I^{0,4} = 2\widehat{\Pi}_{\text{pre},I}^{0,2}\Gamma_I(1-2\Gamma_I) + \widehat{\Pi}_{\text{pre},I}^{0,1}\Gamma_I(1-2\Gamma_I)$$

$$\widehat{\Pi}_I^{1,1} = \widehat{\Pi}_{\text{pre},I}^{1,1}(1-2\Gamma_I)^2$$

$$\widehat{\Pi}_I^{1,2} = \widehat{\Pi}_{\text{pre},I}^{1,2}(1-2\Gamma_I)^2$$

$$\widehat{\Pi}_I^{1,3} = 2\widehat{\Pi}_{\text{pre},I}^{0,1}\Gamma_I(1-2\Gamma_I) + 2\widehat{\Pi}_{\text{pre},I}^{1,1}\Gamma_I(1-2\Gamma_I)$$

$$\widehat{\Pi}_I^{1,4} = 2\widehat{\Pi}_{\text{pre},I}^{1,2}\Gamma_I(1-2\Gamma_I) + 2\widehat{\Pi}_{\text{pre},I}^{1,1}\Gamma_I(1-2\Gamma_I)$$

$$\hat{\Pi}_I^{2,2} = \hat{\Pi}_{\text{pre},I}^{2,2}(1-2\Gamma_I)^2$$

$$\hat{\Pi}_I^{2,3} = 2\hat{\Pi}_{\text{pre},I}^{0,2}\Gamma_I(1-2\Gamma_I) + \hat{\Pi}_{\text{pre},I}^{1,2}\Gamma_I(1-2\Gamma_I)$$

$$\hat{\Pi}_I^{2,4} = 4\hat{\Pi}_{\text{pre},I}^{2,2}\Gamma_I(1-2\Gamma_I) + \hat{\Pi}_{\text{pre},I}^{1,2}\Gamma_I(1-2\Gamma_I)$$

$$\hat{\Pi}_I^{3,3} = 4\hat{\Pi}_{\text{pre},I}^{0,0}\Gamma_I^2 + 2\hat{\Pi}_{\text{pre},I}^{0,2}\Gamma_I^2 + \hat{\Pi}_{\text{pre},I}^{1,1}\Gamma_I^2$$

$$\hat{\Pi}_I^{3,4} = 4\hat{\Pi}_{\text{pre},I}^{0,2}\Gamma_I^2 + 2\hat{\Pi}_{\text{pre},I}^{0,1}\Gamma_I^2 + 2\hat{\Pi}_{\text{pre},I}^{1,2}\Gamma_I^2 + 2\hat{\Pi}_{\text{pre},I}^{1,1}\Gamma_I^2$$

$$\hat{\Pi}_I^{4,4} = 4\hat{\Pi}_{\text{pre},I}^{2,2}\Gamma_I^2 + 2\hat{\Pi}_{\text{pre},I}^{1,2}\Gamma_I^2 + \hat{\Pi}_{\text{pre},I}^{1,1}\Gamma_I^2.$$

#### Text S2: Testing optimization methods

We investigated each optimization method implemented in the optim() function in R (Nelder-Mead, BFGS, CG, L-BFGS-B, and SANN) and found that SANN was the most effective at maximizing the log-likelihood when applying the model to the UK Biobank sequence data. Specifically, we used all five optimization methods (with their default arguments) to fit our model on all 101 MAF intervals and found that SANN produced the highest maximized log-likelihood in 47 MAF intervals, which was the highest count across all of the optimization methods. SANN also produced the highest mean maximized log-likelihood across all 101 MAF intervals.

### Fig S1

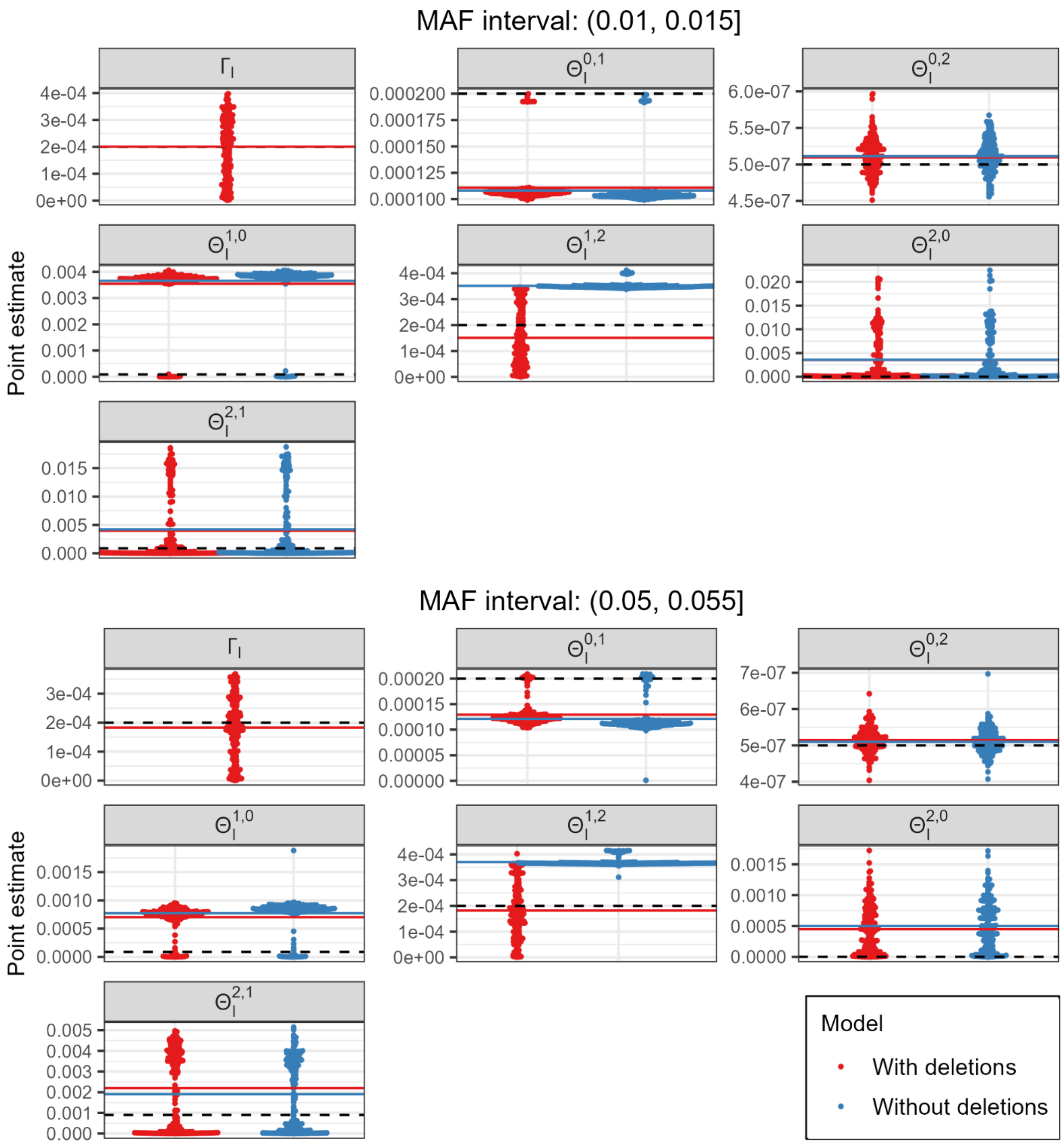


**Fig S1. Distribution of parameter estimates from simulated trio genotype counts from markers in $(0.01, 0.015]$ and $(0.05, 0.055]$.** The dashed black line represents the true parameter value from which the observed trio genotypes are simulated. The red and blue lines represent the sample mean of the estimates from the model with and without deletions respectively.

### Fig S2

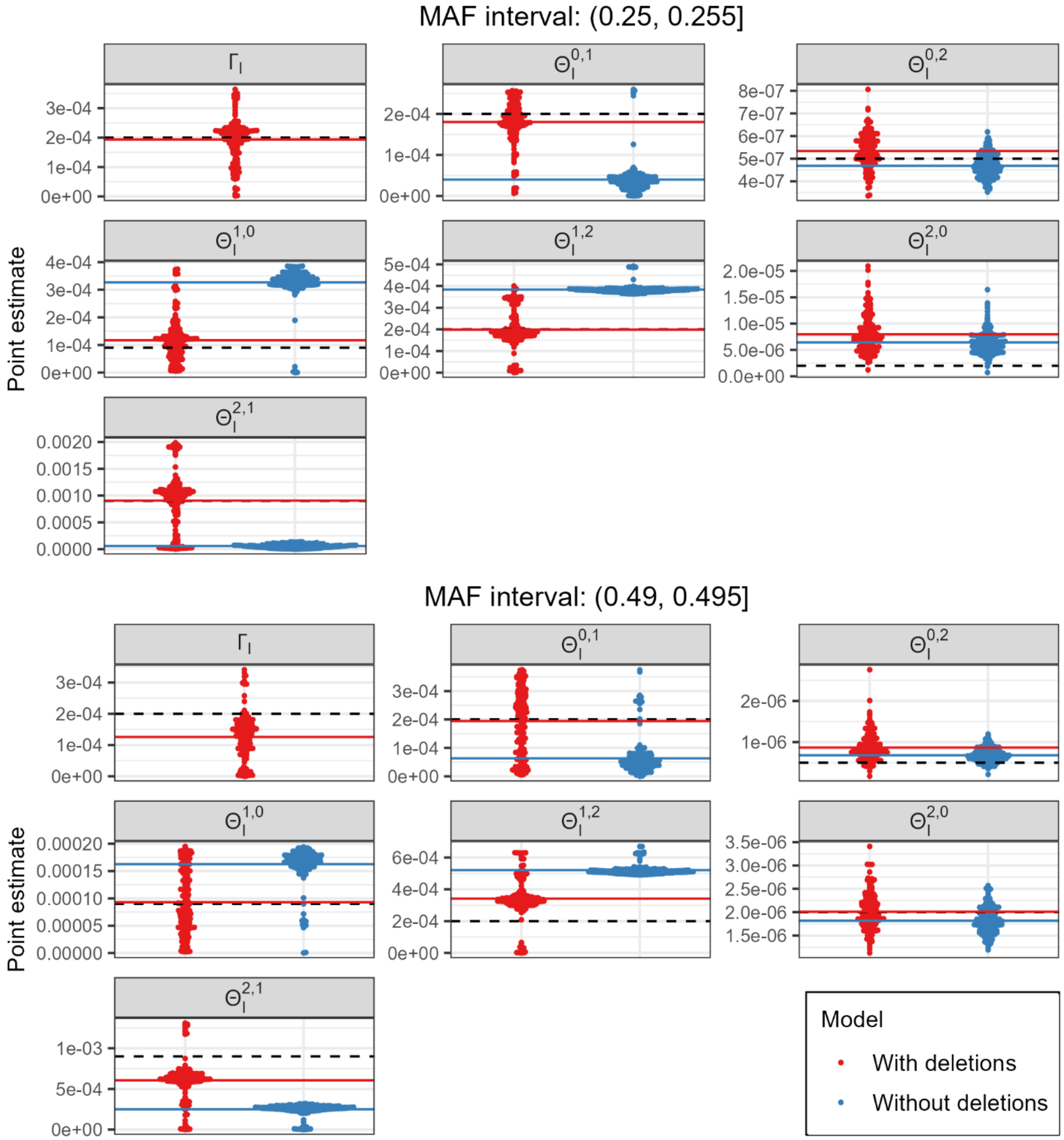


**Fig S2. Distribution of parameter estimates from simulated trio genotype counts from markers in $(0.25, 0.255]$ and $(0.49, 0.495]$.** The dashed black line represents the true parameter value from which the observed trio genotypes are simulated. The red and blue lines represent the sample mean of the estimates from the model with and without deletions respectively.
